# α-Synuclein Strain Propagation is Independent of Cellular Prion Protein Expression in Transgenic Mice

**DOI:** 10.1101/2024.03.27.587028

**Authors:** Raphaella W.L. So, Erica Stuart, Aeen Ebrahim Amini, Adriano Aguzzi, Graham L. Collingridge, Joel C. Watts

## Abstract

The cellular prion protein, PrP^C^, has been postulated to function as a receptor for α-synuclein, potentially facilitating cell-to-cell spreading and/or toxicity of α-synuclein aggregates in neurodegenerative disorders such as Parkinson’s disease. To test this hypothesis, we compared the propagation behavior of two different α-synuclein aggregate strains in M83 transgenic mice that either expressed or did not express PrP^C^. Following intracerebral inoculation with the S or NS strain, the presence of PrP^C^ had minimal influence on α-synuclein strain-specified attributes such as the kinetics of disease progression, the extent of cerebral α-synuclein deposition, selective targeting of specific brain regions and cell types, the morphology of induced α-synuclein deposits, and the structural fingerprints of protease-resistant α-synuclein aggregates. Likewise, there were no appreciable differences in disease manifestation between PrP^C^-expressing and PrP^C^-lacking M83 mice following intraperitoneal inoculation of the S strain. Interestingly, intraperitoneal inoculation with the NS strain resulted in two distinct disease phenotypes, indicative of α-synuclein strain evolution, but this was also independent of PrP^C^ expression. Overall, these results suggest that PrP^C^ plays at most a minor role in the propagation, neuroinvasion, and evolution of α-synuclein strains. Thus, other putative receptors or cell-to-cell propagation mechanisms may play a larger role in the spread of α-synuclein aggregates during disease.

## Introduction

Aggregates of α-synuclein (α-syn) protein are a pathological hallmark of the “synucleinopathies”, neurodegenerative diseases which include Parkinson’s disease (PD) and related disorders such as dementia with Lewy bodies (DLB) and multiple system atrophy (MSA) [1,2]. α-Syn is a 140-amino-acid intrinsically disordered cytosolic protein that localizes to presynaptic nerve terminals under physiological conditions where it is involved in the assembly of SNARE complexes that mediate vesicular membrane fusion [3]. In the synucleinopathies, α-syn becomes phosphorylated at serine-129 (PSyn) and aggregates into β-sheet-rich structural assemblies such as oligomers and fibrils, eventually forming insoluble protein deposits in neurons and glial cells [4,5]. α-Syn plays a pivotal if not causal role in the synucleinopathies, as mutations and copy number variations of the α-syn-coding gene, *SNCA*, have been reported in patients with early-onset PD and DLB [6-10]. Moreover, over-expression of mutant α-syn can lead to neurological symptoms and premature death in transgenic mice [11,12].

Differences in clinical presentation, disease progression rates, and neuropathological features define the various synucleinopathies, and these differences are likely explained by the presence of structurally distinct α-syn aggregates (conformational strains) within the brain in each disease [13-16]. Neuropathologically, the brains of PD and DLB patients have neuronal α-syn inclusions known as Lewy bodies (LBs) and Lewy neurites, while MSA patients exhibit oligodendroglial α-syn deposits termed glial cytoplasmic inclusions. In PD, the appearance of Lewy pathology in the brain appears to follow a hierarchical pattern, with LBs first appearing in the dorsal motor nucleus of the vagus and the olfactory bulb and then spreading to regions of the midbrain and eventually to the cortex as the disease progresses [17]. Along with the observation that LB pathology can spread into tissue grafts [18,19], these findings have led to the hypothesis that α-syn aggregates can propagate from cell to cell within the brain [20,21]. In this “prion-like” model, an α-syn aggregate in one cell undergoes fragmentation to generate smaller seeds, which then escape the cell and are taken up by a neighboring cell where they induce the *de novo* aggregation of normal α-syn. Indeed, α-syn can be transferred between cells [22,23], injection of pre-formed α-syn aggregates into mice induces the aggregation and spreading of α-syn pathology within the brain [24-31], and peripheral injection of mice with α-syn aggregates results in the neuroinvasion of α-syn pathology into the central nervous system [32-37].

As α-syn is an intracellular protein, it remains to be determined how α-syn aggregates might spread from cell to cell across neuronal networks, leading to a cascade of templated protein misfolding. Suggested mechanisms of cellular α-syn transfer include exosomes, endocytosis, receptor-mediated endocytosis, and tunneling nanotubes [38-41]. Putative cell-surface receptors that may mediate the uptake of α-syn oligomers and/or fibrils into cells have been identified, including lymphocyte activation gene 3 (LAG3) [42], the α3 subunit of the sodium-potassium ATPase [43], heparan sulfate proteoglycans [44,45], LRP1 [46], APLP1 [42], neurexin 1β [42,47], and the cellular prion protein (PrP^C^) [48-51].

PrP^C^, encoded by the *PRNP* gene, is a cell-surface neuronal and glial protein more commonly known for its ability to misfold into infectious and pathological protein aggregates that underlie prion diseases in humans and animals such as Creutzfeldt-Jakob disease, scrapie, and chronic wasting disease [52,53]. While its biological role in a non-disease setting is still debated, PrP^C^ has been reported to interact with several pathological protein assemblies, facilitating their spread and/or neurotoxic signaling [54-60]. α-Syn oligomers have been found to interact with regions of the unstructured N-terminal domain of PrP^C^, resulting in neuronal dysfunction via a pathway involving mGluR5, Fyn, and the NMDA receptor [49,55,61]. Upon treatment with α-syn aggregates, cellular and mouse models expressing PrP^C^ internalize more α-syn oligomers or fibrils, exhibit increased α-syn pathology, and/or display more impairment to long-term potentiation than models without PrP^C^ expression or models in which the α-syn binding site on PrP^C^ has been blocked with an antibody [48,50,55]. Furthermore, overexpression of PrP^C^ has been found to exacerbate α-syn aggregation in cells and in mice [50,62]. However, another study reported that α-syn oligomers and PrP^C^ do not interact and that hippocampal neurons from *Prnp* knockout (*Prnp*^0/0^) mice were no less susceptible to α-syn aggregate-induced toxicity than neurons from wild-type mice [63]. Thus, the role of PrP^C^ in the propagation of α-syn aggregates remains to be completely resolved.

In this study, we used M83 transgenic mice expressing mutant human α-syn to investigate the role of PrP^C^ in the propagation and neuroinvasion of two distinct conformational strains of α-syn aggregates [12,31]. We hypothesized that if PrP^C^ plays a significant role in the propagation of pathogenic α-syn seeds, M83 mice that lack PrP^C^ expression should succumb to neurological disease later, or not at all, upon injection with α-syn aggregates, leading to fewer PSyn inclusions in the brain. However, M83 mice inoculated intracerebrally or intraperitoneally with α-syn aggregates developed PSyn inclusions and motor dysfunction regardless of *Prnp* genotype. Moreover, the presence or absence of PrP^C^ had no discernible effect on the neuropathological or biochemical attributes of the S and NS strains. We observed subtle, but inconsistent effects of PrP^C^ expression on disease kinetics following either intracerebral or intraperitoneal inoculation with the NS strain, whereas no such differences were observed with the S strain. Thus, we conclude that PrP^C^ has minimal influence on the propagation and neuroinvasion of α-syn strains in a transgenic mouse model of synucleinopathy.

## Materials and Methods

### Mice

Homozygous M83 mice (stock number: 004479), which are on a mixed C57BL/6 and C3H background [12], and non-transgenic B6C3F1 mice (stock number: 100010) were purchased from The Jackson Lab. These mice were intercrossed to generate hemizygous M83 mice, which were then crossed with the ZH3 strain of *Prnp*^0/0^ mice, which are on a co-isogenic C57BL/6 background [64]. *Prnp*^0/+^ mice that were hemizygous for the M83 transgene were then intercrossed with non-transgenic C57BL/6 mice and ZH3 *Prnp*^0/0^ mice, respectively, to generate the M83-*Prnp*^+/+^ and M83-*Prnp*^0/0^ lines. All animal experiments were conducted in accordance with a protocol (AUP #4263.19) approved by the University Health Network Animal Care Committee. Mice were given free access to food and water and were maintained on a 12 h light/12 h dark cycle. Both male and female mice were used for all experiments.

### Inoculations

Brains of clinically ill mice from the third passage of the S or NS strains in hemizygous M83 mice were used as inocula [31]. Brain homogenates [10% (w/v)] in Dulbecco’s PBS (ThermoFisher Scientific #14190144) were generated using CK14 soft tissue homogenizing tubes containing 1.4 mm zirconium oxide beads (Bertin Technologies #P000912-LYSK0-A) and a Minilys homogenizer (Bertin Technologies), then aliquoted and stored at-80 °C. Prior to inoculation, brain homogenates were diluted to 1% (w/v) using 5% (w/v) BSA prepared in PBS. For intracerebral inoculations, M83-*Prnp*^+/+^ and M83-*Prnp*^0/0^ mice at 5 weeks of age were injected non-stereotactically into the right cerebral hemisphere at a depth of ∼3 mm using a tuberculin syringe with an attached 27 gauge, 0.5-inch needle (BD Biosciences #305945). Each mouse received 30 µL of diluted brain homogenate. For the intraperitoneal inoculations, M83-*Prnp*^+/+^ and M83-*Prnp*^0/0^ mice at 5-6 weeks of age were injected into the peritoneal cavity with 200 µL of diluted brain homogenate. Inoculated mice and uninoculated controls were monitored biweekly for the development of neurological symptoms such as hindlimb paralysis or tremor, bradykinesia, loss of grip strength, and weight loss [31]. Observers were blind to the inoculum but not the mouse genotype. Mice were euthanized once advanced hindlimb paralysis that affected their ability to ambulate was noted and were then perfused with saline and their brains removed and bisected parasagittally. The left hemisphere was flash-frozen on dry ice and stored at-80°C and the right hemisphere was fixed in 10% (v/v) neutral buffered formalin. Mice found dead in their cages or that were euthanized due to intercurrent or atypical (non-synucleinopathy) illness were excluded from the study (**Table S1**). Synucleinopathy status was assessed by examining the presence of protease-resistant α-syn in brain homogenates, as described below.

### Detergent insolubility assays

Brain homogenates from uninoculated or inoculated mice were generated as described above. Nine volumes of brain homogenate were mixed with one volume of 10X detergent buffer [5% (w/v) sodium deoxycholate and 5% (v/v) Nonidet P-40 in PBS] containing Pierce Universal Nuclease (ThermoFisher Scientific #88701) and Halt Phosphatase Inhibitor (ThermoFisher Scientific #78420), and incubated on ice for 20 min. Brain extracts were clarified by centrifugation at 1,500x *g* for 5 min at 4 °C and then protein concentrations in the supernatant were determined using the bicinchoninic acid (BCA) assay (ThermoFisher Scientific #23227). Detergent-extracted brain homogenates were diluted to a concentration of 5 mg/mL in 1X detergent buffer and then incubated on ice for 20 min. The samples were then ultracentrifuged in a TLA-55 rotor (Beckman Coulter) at 100,000x *g*. for 1 h at 4 °C. The pellets were resuspended in loading buffer [1X Bolt LDS sample buffer (ThermoFisher Scientific #B0007) containing 2.5% (v/v) β-mercaptoethanol] then heated at 95 °C for 10 min before storage at-20 °C or analysis by immunoblotting. For quantification of detergent-insoluble α-syn levels, band intensity levels were determined by densitometry using ImageJ.

### Protease digestion assays

Protein concentrations in detergent-extracted brain homogenates were measured using the BCA assay. For thermolysin (TL) digestions (Sigma-Aldrich #T7902), brain extract at a concentration of 5 mg/mL was treated with 50 µg/mL protease (TL:protein ratio of 1:100) with shaking at 600 r.p.m. for 1 h, and then the reaction was stopped by adding EDTA to a final concentration of 5 mM. For proteinase K (PK) digestions (ThermoFisher Scientific #EO0491), a protease concentration of 100 µg/mL was used instead (PK:protein ratio of 1:50), and the reaction was stopped by adding PMSF to a final concentration of 4 mM. Protease-digested samples were then ultracentrifuged in a TLA-55 rotor at 100,000x *g*. for 1 h at 4 °C. The pellets were resuspended in loading buffer and then heated at 95 °C for 10 min before analysis by immunoblotting.

### Immunoblotting

Samples prepared in loading buffer were run on 12% Bolt Bis-Tris Plus mini gels (ThermoFisher Scientific #NW00120BOX or NW00122BOX) at 165 V for ∼50 min. Contents of the gel were transferred onto 0.45 μm Immobilon-P PVDF membranes (MilliporeSigma #IPVH00010) in transfer buffer [25 mM Tris-HCl pH 8.3, 0.192 M glycine, and 20% (v/v) methanol] at 35 V for 70 min. The proteins were crosslinked to the membrane using 0.4% (v/v) paraformaldehyde (Electron Microscopy Services #15711) in PBS for 30 min at 22 °C [65], and then rinsed twice with TBS containing 0.1% (v/v) Tween-20 (TBST). Membranes were treated with blocking buffer [5% (w/v) skim milk in TBST] for 1 h at 22 °C and then incubated with primary antibody (diluted in blocking buffer) overnight at 4 °C. The following primary antibodies were used in this study: anti-total α-syn mouse monoclonal Syn-1 (1:10,000 dilution; BD Biosciences #610786), anti-PSyn rabbit monoclonal EP1536Y (1:10,000 dilution; Abcam #ab51253), anti-human α-syn rabbit monoclonal MJFR1 (1:10,000 dilution; Abcam #ab138501), anti-mouse α-syn rabbit monoclonal D37A6 (1:1,000 dilution; Cell Signaling #4179), and humanized recombinant anti-PrP^C^ Fab HuM-D18 (1:5,000 dilution) [66]. Membranes were washed three times with TBST for 10 min, then incubated in blocking buffer containing horseradish peroxidase-conjugated secondary antibodies at 1:10,000 dilution (Bio-Rad #172-1011 or 172-1019, or ThermoFisher Scientific #31414) for 1 h at 22 °C. After three washes in TBST, the blots were developed using Western Lightning enhanced chemiluminescence Pro (Revvity #NEL122001EA) or SuperSignal West Dura Extended Duration Substrate (ThermoFisher Scientific #37071) and were either exposed to HyBlot CL X-ray film (Thomas Scientific #1141J52) or imaged using the LiCor Odyssey Fc. For assessment of equal protein loading, membranes were reprobed with the anti-actin 20-33 rabbit polyclonal antibody (1:10,000 dilution; Sigma Aldrich #A5060). For comparison of protease-resistant α-syn banding patterns, the amount of protein loaded was empirically adjusted so the bands were of similar signal strength.

### Human α-syn ELISA

Protein concentrations in detergent-extracted brain homogenates were measured using the BCA assay, then samples were diluted to a concentration of 5 µg/mL using PBS. The concentrations of human α-synuclein were measured using a human α-syn-specific ELISA (ThermoFisher Scientific #KHB0061) according to the manufacturer’s protocol. Data points were averaged across two independent trials.

### Immunohistochemistry

Perfused half brains were fixed in formalin and then processed, embedded in paraffin, and cut into 4 µm sagittal sections and adhered to Superfrost Plus glass slides (Fisher Scientific #22037246). Slides were heated at 60 °C for 20 min for deparaffinization using xylenes and rehydrated using a graded series of ethanol. Epitope retrieval was performed by submersion in citrate buffer [10 mM sodium citrate pH 6, 0.05% (v/v) Tween-20] and steaming for 10 min in an Instant Pot under high pressure. After the slides were cooled to 22 °C, they were washed in running tap water for 10 min. Endogenous peroxidase activity was quenched by incubating the slides in 3% H2O2 prepared in PBS for 5 min then washing three times with PBS containing 0.1% (v/v) Tween-20 (PBST), also for 5 min each. Slides were blocked with 2.5% (v/v) normal horse serum (Vector Laboratories #MP-7401) for 1 h at 22 °C. Primary antibody incubation was performed overnight at 4 °C with rabbit monoclonal anti-PSyn EP1536Y diluted 1:320,000 in DAKO antibody diluent (Agilent #S0809). After three 5-min washes with PBST, slides were incubated with ImmPress horseradish-peroxidase-labeled horse anti-rabbit detection kit (Vector Laboratories #MP-7401) at 22 °C for 30 min, then rinsed with PBST followed by three 5-min PBST washes. Slides were developed with ImmPACT 3,3’-diaminobenzidine peroxidase substrate (Vector Laboratories #SK-4105) for 1 min, rinsed under running tap water for 10 min, counterstained with hematoxylin (Sigma-Aldrich #GHS132) for 2 min, then mounted with Cytoseal 60 (Epredia #8310-4). Tissue slides were scanned using the TissueSnap and TissueScope LE120 system (Huron Digital Pathology) and then viewed using PMA.start (Pathomation). The percentage area covered by PSyn deposits in the pons, midbrain, and hypothalamus was quantified using ImageJ as previously described [67], except that a threshold range of 0-75 was used. The number of PSyn-positive neurons in the cerebral cortex and the CA1 region of the hippocampus, as well as the number of PSyn-positive astrocytes in the thalamus were manually counted.

### Statistical analysis

All statistical analysis was performed using GraphPad Prism (version 10.2.1) with a significance threshold of P = 0.05. Survival curves for inoculated mice were compared using the Log-rank test. It was not assumed that experimental data was normally distributed, and therefore non-parametric tests were used to assess statistical significance. For comparisons involving two samples, a Mann-Whitney test was used. For comparisons involving three or more groups, a Kruskal-Wallis test followed by Dunn’s multiple comparisons test was utilized. For the analysis of α-syn deposition in regions of the midbrain as well as the morphology of induced α-syn deposits, a two-way ANOVA followed by Šídák’s multiple comparisons test was performed.

## Results

### Generation of M83-Prnp^+/+^ and M83-Prnp^0/0^ mice

To test whether PrP^C^ plays a role in facilitating the propagation of α-syn aggregates, we used the M83 transgenic mouse model, which expresses human α-syn with the A53T mutation linked to an early-onset genetic form of PD [12]. Hemizygous M83 mice (M83^+/-^) do not spontaneously develop a synucleinopathy up to 600 days of age, but develop progressive signs of motor dysfunction accompanied by cerebral deposition of aggregated and phosphorylated α-syn if injected with a pre-existing source of α-syn aggregates [26,27,31]. We crossbred M83^+/-^ mice with *Prnp*^0/0^ mice on a C57BL/6 background [64] and then backcrossed to wild-type C57BL/6 or *Prnp*^0/0^ mice to create M83^+/-^ mice that either express or do not express PrP^C^ (M83-*Prnp*^+/+^ and M83-*Prnp*^0/0^ mice, respectively) with otherwise minimal genetic differences. As expected, brain extracts from the M83-*Prnp*^0/0^ mice did not exhibit any PrP^C^ (**Fig. 1a**), and there were no differences in the levels of mouse, human, or total (mouse + human) α-syn between the M83-*Prnp*^+/+^ and M83-*Prnp*^0/0^ lines (**Fig. 1b**). The lack of a difference in human α-syn levels between M83-*Prnp*^+/+^ and M83-*Prnp*^0/0^ mice was further confirmed by ELISA (**Fig. 1c**). In the absence of inoculation, both M83-*Prnp*^+/+^ and M83-*Prnp*^0/0^ mice remained free of neurological dysfunction up to 18 months of age (**Table 1**).

**Figure 1.**
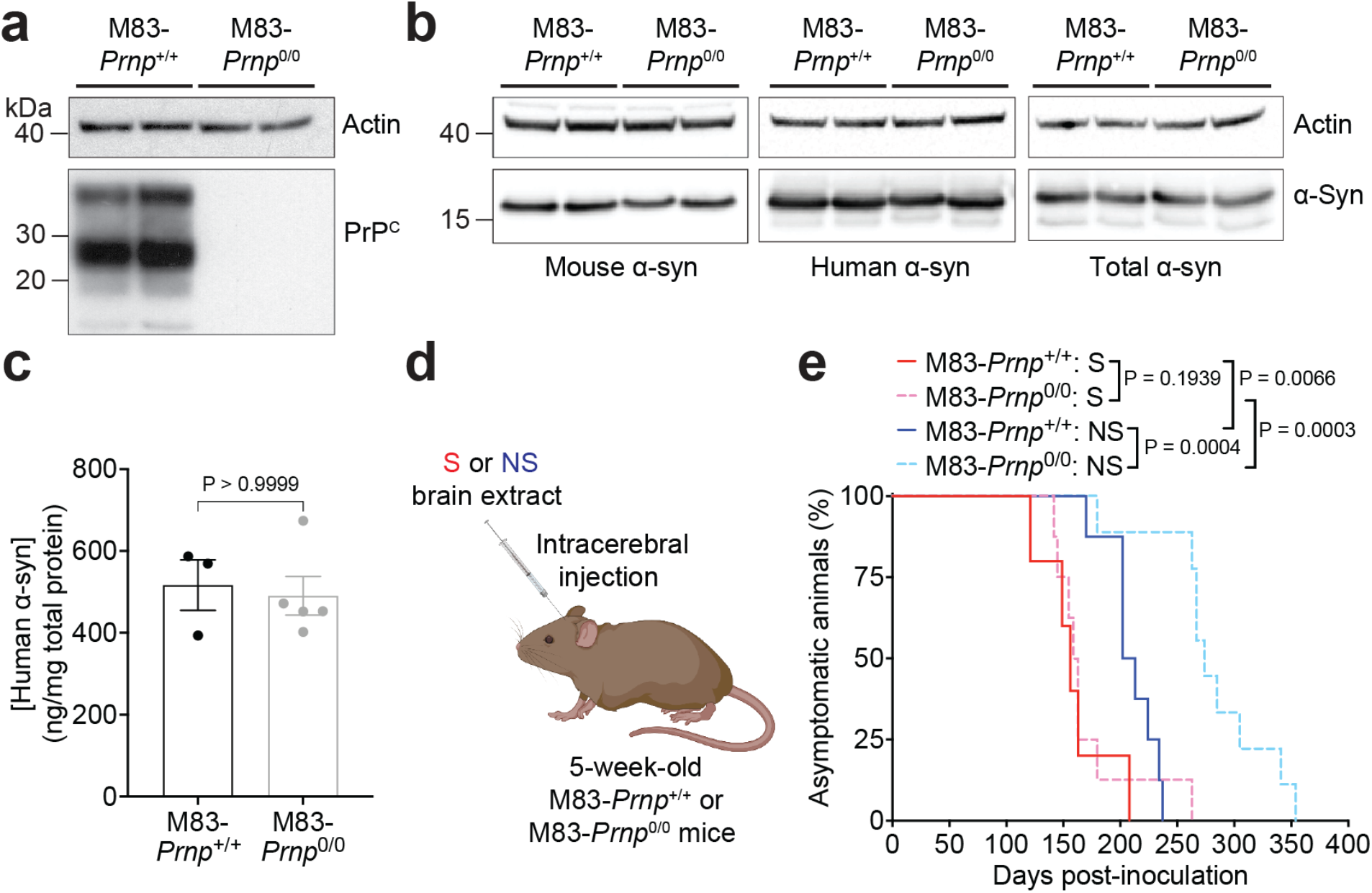
Intracerebral inoculation of M83-*Prnp*^+/+^ and M83-*Prnp*^0/0^ mice with the S and NS strains. a) Immunoblot for PrP^C^ in brain extracts from 18-month-old uninoculated M83-*Prnp*^+/+^ and M83-*Prnp*^0/0^ mice (2 independent mice per line). The blot was reprobed with an actin antibody. **b)** Immunoblots for mouse α-syn, human α-syn, and total (mouse + human) α-syn in brain extracts from 18-month-old uninoculated M83-*Prnp*^+/+^ and M83-*Prnp*^0/0^ mice (2 independent mice per line). Each blot was reprobed with an actin antibody. **c)** ELISA results for human α-syn in brain extracts from 18-month-old uninoculated M83-*Prnp*^+/+^ (n = 3) and M83-*Prnp*^0/0^ (n = 5) mice. The graph displays mean ± s.e.m. Statistical significance was assessed using a Mann-Whitney test. **d)** Schematic of intracerebral inoculation experiments in M83-*Prnp*^+/+^ and M83-*Prnp*^0/0^ mice. **e)** Kaplan-Meier survival curves for M83-*Prnp*^+/+^ and M83-*Prnp*^0/0^ mice inoculated intracerebrally with S or NS strain (n = 5-9 mice per experimental condition). Statistical significance was assessed using the Log-rank test.

**Table 1.** Disease incubation periods in inoculated M83-*Prnp*^+/+^ and M83-*Prnp*^0/0^ mice.

| Inoculum | Inoculation route | *Prnp* genotype | Incubation period (days ± s.e.m.) | Symptomatic mice | TL-resistant α-syn |
| --- | --- | --- | --- | --- | --- |
| None | N/A | +/+ | >500<sup>1</sup> | 0/3 | 0/3 |
|  | N/A | 0/0 | >500<sup>1</sup> | 0/5 | 0/5 |
| S | IC | +/+ | 159 ± 14 | 5/5 | 5/5 |
| | IC | 0/0 | 171 ± 14 | 8/8 | 8/8 |
| | IP | +/+ | 227 ± 17 | 10/10 | 10/10 |
| | IP | 0/0 | 252 ± 12 | 5/5 | 5/5 |
| NS | IC | +/+ | 210 ± 8 | 8/8 | 8/8 |
| | IC | 0/0 | 282 ± 17 | 9/9 | 9/9 |
| | IP | +/+ | 389 ± 40 | 5/6<sup>2</sup> | 5/6 |
| | IP | 0/0 | 293 ± 35 | 6/6 | 6/6 |

<sup>1</sup>Asymptomatic uninoculated M83-*Prnp*<sup>+/+</sup> and M83-*Prnp*<sup>0/0</sup> mice were collected at 18 months of age (~500 days following mock inoculation)
<sup>2</sup>One M83-*Prnp*<sup>+/+</sup> mouse injected IP with NS remained asymptomatic at 540 days post-inoculation

### Transmission of S and NS strains to mice by intracerebral inoculation

We hypothesized that if PrP^C^ facilitates the pathological spread of α-syn aggregates, M83-*Prnp*^+/+^ mice should develop neurological disease faster than M83-*Prnp*^0/0^ mice following inoculation with α-syn aggregates, or that there should be more cerebral α-syn deposition in the former compared to the latter. Previously, we described two distinct α-syn conformational strains made from recombinant human α-syn with the A53T mutation [31]. When intracerebrally injected into M83^+/-^ mice, fibrils made in the presence of salt (S fibrils) caused faster neurological disease than those made in the absence of salt (NS fibrils). Serial passaging of brain extracts from mice originally injected with S or NS fibrils led to the generation of the S and NS strains of α-syn aggregates. The S and NS strains produce distinct disease phenotypes in M83^+/-^ mice, characterized by differences in disease kinetics, the morphology of cerebral α-syn inclusions, the brain regions and cell-types targeted, as well the biochemical properties of the induced α-syn aggregates [31].

M83-*Prnp*^+/+^ and M83-*Prnp*^0/0^ mice underwent intracerebral (IC) inoculation with M83^+/-^ mouse brain extract containing either the S or NS strain of α-syn aggregates and were then monitored longitudinally for the development of progressive motor impairment (**Fig. 1d**). All M83-*Prnp*^+/+^ and M83-*Prnp*^0/0^ mice injected intracerebrally with either the S or NS strain developed neurological disease (**Fig. 1e**, **Table 1**). For the S strain, there was no significant difference in the disease kinetics for the M83-*Prnp*^+/+^ and M83-*Prnp*^0/0^ lines (**Fig. 1e**). M83-*Prnp*^+/+^ mice inoculated with the NS strain succumbed to disease modestly, but significantly earlier than M83-*Prnp*^0/0^ mice. Consistent with our previous results [31], the S strain induced neurological disease earlier than the NS strain following IC inoculation in both the M83-*Prnp*^+/+^ and M83-*Prnp*^0/0^ lines. Mice that were euthanized due to non-synucleinopathy illness were excluded from the study, and this occurred in both the M83-*Prnp*^+/+^ and M83-*Prnp*^0/0^ lines (**Table S1**).

*Biochemical characterization of induced α-syn aggregates following intracerebral inoculation* In brain extracts prepared from aged uninoculated M83-*Prnp*^+/+^ and M83-*Prnp*^0/0^ mice, detergent-insoluble PSyn, the disease-associated species, was not detected (**Fig. 2a**). In contrast, brain extracts from symptomatic S and NS strain-inoculated M83-*Prnp*^+/+^ and M83-*Prnp*^0/0^ mice exhibited abundant PSyn. Detergent-insoluble total α-syn, which includes both phosphorylated and non-phosphorylated species, was invariably present in all brain extracts, regardless of symptomatic status, likely due to overexpression of A53T-mutant human α-syn in M83^+/-^ mice. For both the S and NS strains, there were no significant differences in the relative amounts of detergent-insoluble PSyn between the M83-*Prnp*^+/+^ and M83-*Prnp*^0/0^ lines (**Fig. 2b**). However, mice inoculated with the NS strain exhibited ∼4-fold higher levels of detergent-insoluble PSyn than mice injected with the S strain (**Fig. 2a, c**), probably because the NS strain accumulates in a greater number of brain regions [31]. Protease digestion assays were also performed, as protease resistance is another indicator of disease-associated α-syn aggregates. As expected, brain extracts from uninoculated asymptomatic M83-*Prnp*^+/+^ and M83-*Prnp*^0/0^ mice did not contain insoluble protease-resistant α-syn (**Fig. 2d**). Meanwhile, S and NS-inoculated mice had insoluble α-syn resistant to both thermolysin (TL) and proteinase K (PK) digestion.

**Figure 2.**
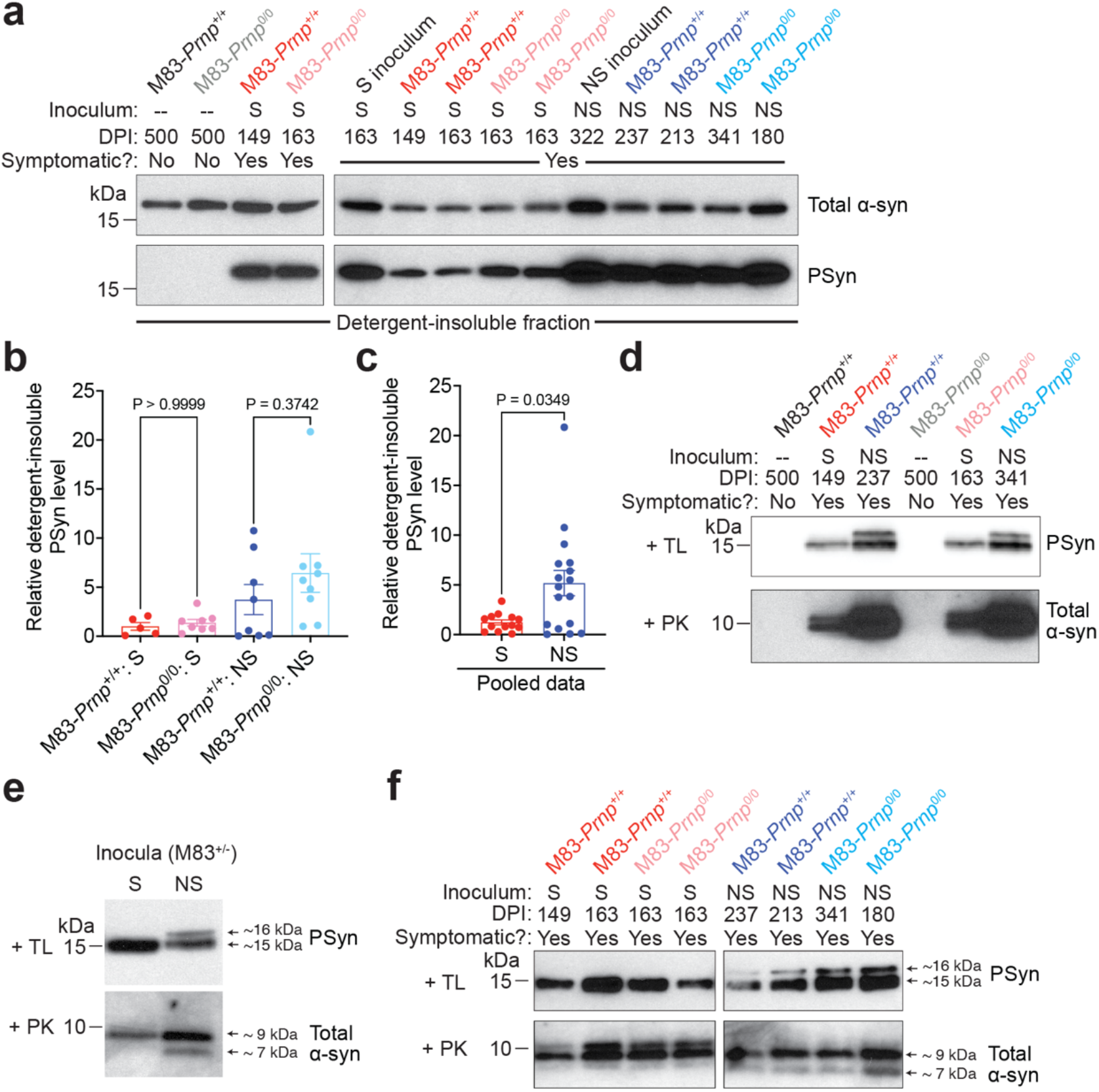
PrP^C^ does not affect the biochemical attributes of the S and NS strains following intracerebral inoculation of M83 mice. a) Immunoblots of detergent-insoluble α-syn in brain extracts from symptomatic M83-*Prnp*^+/+^ and M83-*Prnp*^0/0^ mice at the indicated days post-inoculation (DPI) with S or NS strain compared with 18-month-old uninoculated (--) mice, probed for total α-syn and PSyn. In the left blots, one mouse is shown for each experimental condition. In the right blots, two distinct mice per experimental condition are shown, along with their respective inocula. **b)** Quantification of detergent-insoluble PSyn levels in brain extracts from the 4 experimental groups (n = 5-9 mice per experimental condition). The graph displays mean ± s.e.m. Statistical significance was assessed using a Kruskal-Wallis test followed by Dunn’s multiple comparisons test. **c)** Pooled detergent-insoluble PSyn data for M83-*Prnp*^+/+^ and M83-*Prnp*^0/0^ mice inoculated with either S (n = 13) or NS (n = 17) strain. The graph displays mean ± s.e.m. Statistical significance was assessed using a Mann-Whitney test. **d)** Immunoblots of detergent-insoluble PSyn in brain extracts from uninoculated, S-inoculated, and NS-inoculated M83-*Prnp*^+/+^ and M83-*Prnp*^0/0^ mice after thermolysin (TL) digestion, as well as detergent-insoluble total α-syn after proteinase K (PK) digestion. **e)** Immunoblots for detergent-insoluble TL-resistant PSyn and PK-resistant total α-syn in the S and NS inocula used for the intracerebral inoculations. **f)** Immunoblots for detergent-insoluble TL-resistant PSyn and PK-resistant total α-syn in brain extracts from symptomatic S-or NS-inoculated M83-*Prnp*^+/+^ and M83-*Prnp*^0/0^ mice at the indicated DPI. Two distinct mice are shown per experimental condition.

The banding patterns of protease-resistant α-syn fragments following digestion with TL or PK can be used to infer structural differences between α-syn aggregates [31,68]. After digestion with TL, a single ∼15 kDa protease-resistant PSyn band is observed for the S strain, whereas an additional band with a slightly higher molecular weight is observed for the NS strain (**Fig. 2e**). Similarly, following PK digestion and detection with an antibody against total α-syn, a prominent ∼9 kDa band is present for the S strain, while a second band at ∼7 kDa is present for the NS strain. We therefore compared the banding patterns of protease-resistant α-syn fragments between M83-*Prnp*^+/+^ and M83-*Prnp*^0/0^ mice inoculated with either the S or NS strains. For either strain, the *Prnp* genotype of the M83 mice had no influence on the banding pattern of the induced α-syn aggregates following protease digestion (**Fig. 2f**). Thus, there was no detectable conformational difference in the two strains of α-syn aggregates propagated in M83-*Prnp*^+/+^ versus M83-*Prnp*^0/0^ mice.

### Neuropathological characterization of induced α-syn aggregates following intracerebral inoculation

Both the S and NS strains produce robust PSyn deposition in regions of the hindbrain in M83^+/-^ mice [31]. Thus, we compared the relative amounts of PSyn deposition in the pons, midbrain, and hypothalamus of M83-*Prnp*^+/+^ and M83-*Prnp*^0/0^ mice following intracerebral inoculation with the S or NS strain. For both α-syn strains, there was no significant difference in the extent of PSyn deposition in these three brain regions between M83-*Prnp*^+/+^ and M83-*Prnp*^0/0^ mice (**Fig. 3a, b**). As expected, the levels of PSyn deposition were significantly higher in the brains of mice inoculated with the S or NS strain compared to aged uninoculated mice, which did not exhibit any PSyn pathology (**Fig. 3a, c**). Moreover, PSyn deposition levels in the three hindbrain regions did not significantly differ between the S and NS strains (**Fig. S1**). Since the S and NS strains differentially target other brain regions and cell types in M83^+/-^ mice [31], we also examined whether the presence or absence of PrP^C^ had any effect on α-syn strain-specific selective vulnerability. The NS strain preferentially induced PSyn deposition in the hippocampus, but this was not dependent on whether PrP^C^ was expressed in the brain (**Fig. 4a-c**). Similarly, the ability of the NS strain to preferentially target cortical neurons was unaltered between M83-*Prnp*^+/+^ and M83-*Prnp*^0/0^ mice (**Fig. 4d-f**). While both the S and NS strain predominantly form PSyn deposits in neurons, PSyn inclusions in astrocytes are also observed in M83^+/-^ mice inoculated with the NS strain [31]. However, the number of PSyn-positive astrocytes in the thalamus of animals inoculated intracerebrally with the NS strain did not significantly differ between M83-*Prnp*^+/+^ and M83-*Prnp*^0/0^ mice (**Fig. 4g-i**). Finally, since the morphology of induced PSyn deposits in the midbrain of M83^+/-^ mice differs for the S and NS strain [31], we compared the morphology of the PSyn deposits in intracerebrally inoculated M83-*Prnp*^+/+^ and M83-*Prnp*^0/0^ mice. Consistent with our previous findings, neurons in the midbrain of mice inoculated with the S strain exhibited the expected “ring-like” morphology whereas neurons in NS-inoculated mice displayed a “LB-like” morphology (**Fig. 5a**). No significant differences in the morphologies of induced PSyn deposits were found between M83-*Prnp*^+/+^ and M83-*Prnp*^0/0^ mice inoculated intracerebrally with the S or NS strains (**Fig. 5b**). Thus, the presence or absence of PrP^C^ has no discernible effect on the neuropathological manifestation of the S or NS α-syn conformational strains.

**Figure 3.**
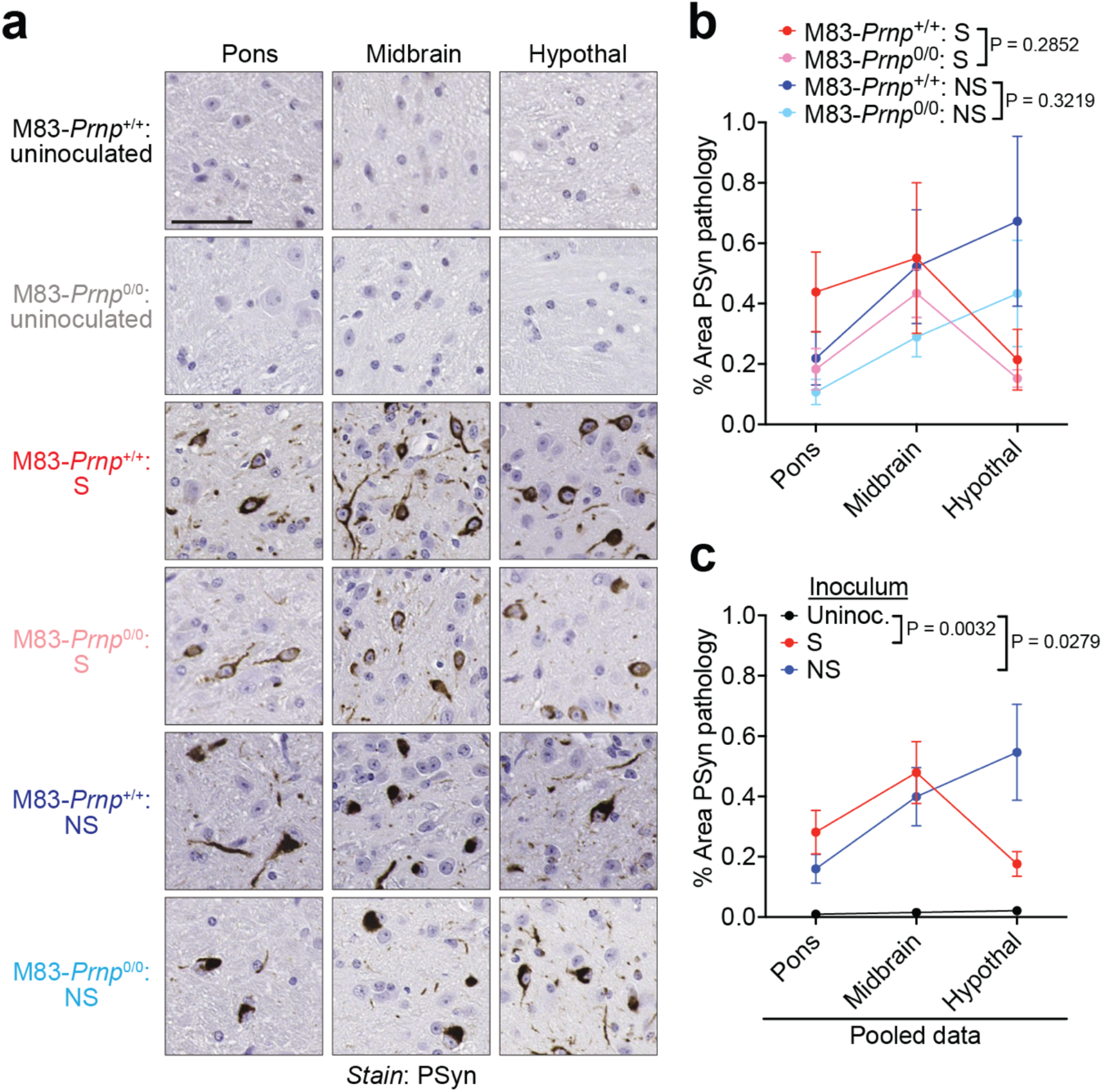
PrP^C^ does not affect the cerebral deposition of α-syn aggregates following intracerebral inoculation of M83 mice with the S and NS strains. a) Representative images of PSyn-stained sections from the pons, midbrain, and hypothalamus of 18-month-old uninoculated M83-*Prnp*^+/+^ and M83-*Prnp*^0/0^ mice as well as symptomatic mice inoculated intracerebrally with the S or NS strains. Scale bar = 50 µm. **b)** Quantification of the area covered by PSyn staining in sections from the indicated brain regions of symptomatic M83-*Prnp*^+/+^ and M83-*Prnp*^0/0^ mice inoculated with either S or NS (n = 5-9 mice per experimental condition). The graph displays mean ± s.e.m. Statistical significance was assessed using two-way ANOVA. **c)** Pooled quantitative PSyn staining data for M83-*Prnp*^+/+^ and M83-*Prnp*^0/0^ mice inoculated with either S (n = 13) or NS (n = 17) strain. Data from uninoculated mice (n = 7) is also shown. The graph displays mean ± s.e.m. Statistical significance was assessed using two-way ANOVA.

**Figure 4.**
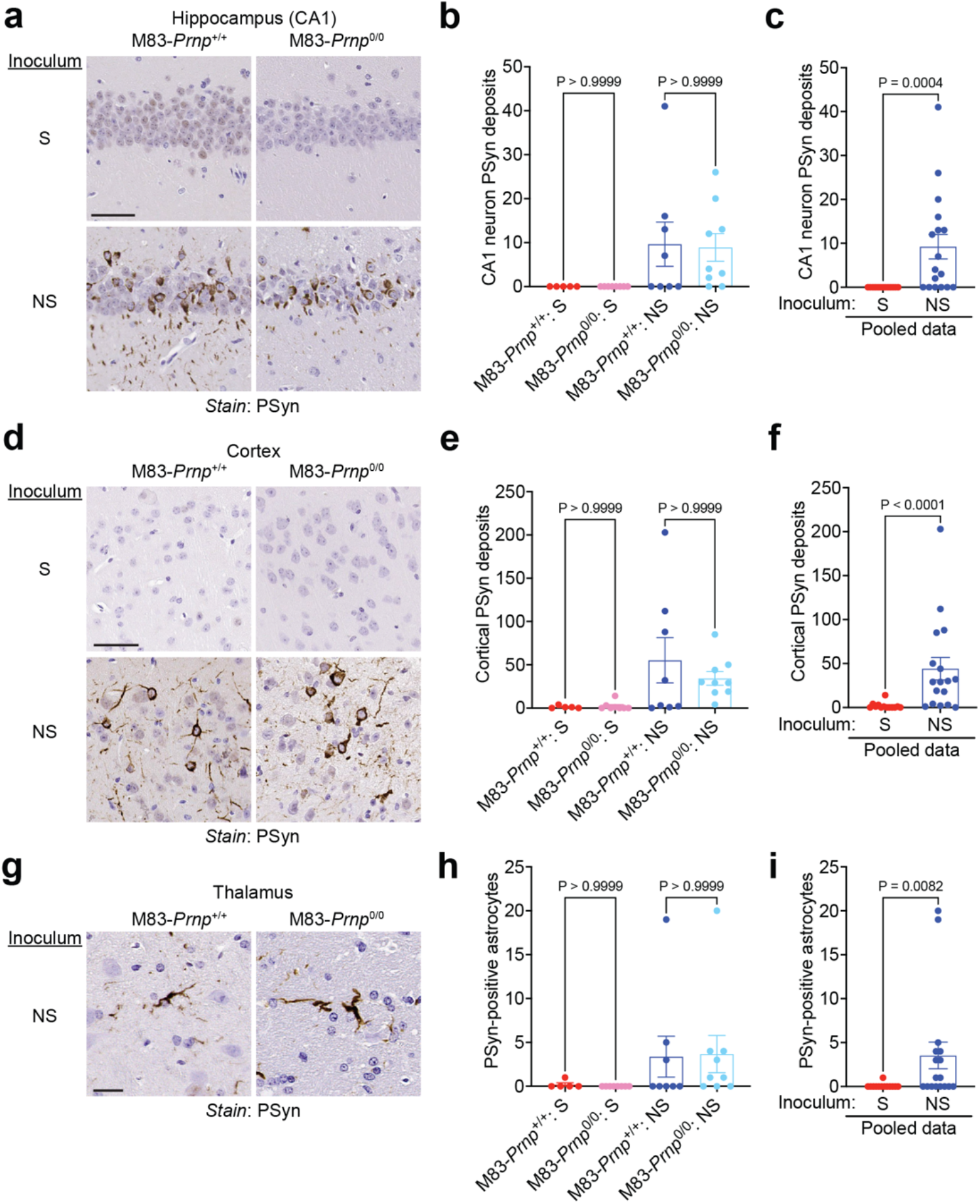
PrP^C^ does not affect the selective targeting of distinct brain regions and cell types by α-syn strains in M83 mice. a, d) Representative images of PSyn-stained sections from the hippocampal CA1 region (**a**) or cortex (**d**) from symptomatic M83-*Prnp*^+/+^ and M83-*Prnp*^0/0^ mice inoculated intracerebrally with the S or NS strains. Scale bar = 50 µm. **b, e)** Quantification of the number of PSyn-positive neurons in the hippocampal CA1 region (**b**) or cortex (**e**) of inoculated mice (n = 5-9 mice per experimental condition). **c, f)** Pooled PSyn-positive neuronal counts in the hippocampal CA1 region (**c**) or cortex (**f**) of M83-*Prnp*^+/+^ and M83-*Prnp*^0/0^ mice inoculated with either S (n = 13) or NS (n = 17) strain. **g)** Representative images of PSyn-stained sections from the thalamus of symptomatic M83-*Prnp*^+/+^ and M83-*Prnp*^0/0^ mice inoculated with the NS strain. Scale bar = 20 µm. **h)** Quantification of the number of PSyn-positive astrocytes in the thalamus of S-or NS-inoculated mice (n = 5-9 mice per experimental condition). **i)** Pooled PSyn-positive astrocyte counts in the thalamus of M83-*Prnp*^+/+^ and M83-*Prnp*^0/0^ mice inoculated with either S (n = 13) or NS (n = 17) strain. All graphs display mean ± s.e.m. In panels b, e, and h, statistical significance was assessed using a Kruskal-Wallis test followed by Dunn’s multiple comparisons test. In panels c, f, and i, statistical significance was assessed using a Mann-Whitney test.

**Figure 5.**
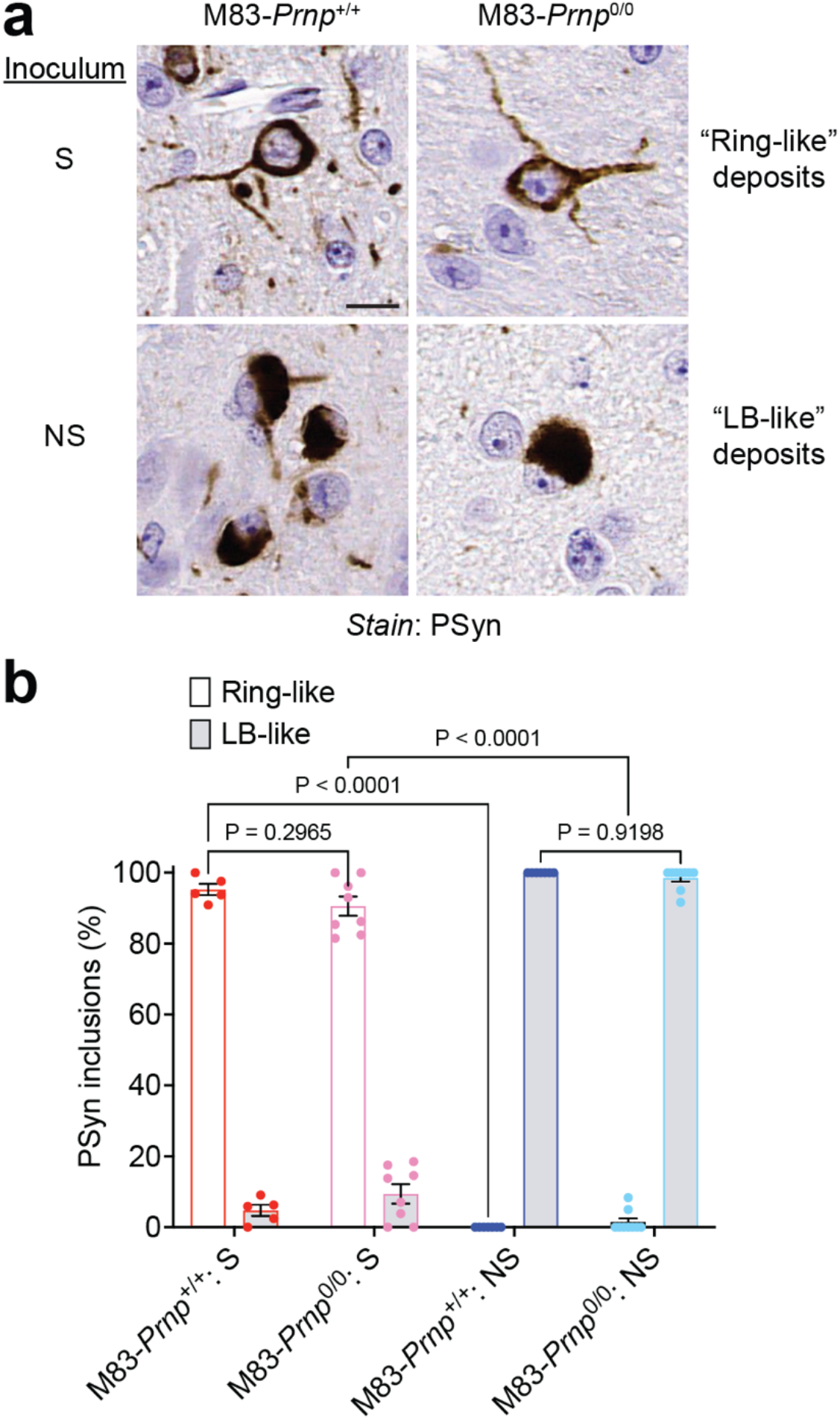
PrP^C^ does not affect α-syn strain-specified differences in the morphology of induced α-syn aggregates in M83 mice. a) Representative images of PSyn-stained sections from the midbrain of symptomatic M83-*Prnp*^+/+^ and M83-*Prnp*^0/0^ mice inoculated intracerebrally with the S or NS strains. Scale bar = 10 µm. **b**) Quantification of the relative percentages of ring-like and LB-like PSyn inclusions in the midbrain of S-or NS-inoculated mice (n = 5-9 mice per experimental condition). The graph displays mean ± s.e.m. Statistical significance was assessed using two-way ANOVA.

### Transmission of S and NS strains to mice by intraperitoneal inoculation

Peripheral administration of α-syn seeds in M83 mice can lead to motor dysfunction and cerebral α-syn deposition, suggesting that α-syn aggregates can propagate from the periphery to the central nervous system [34-37,69,70]. Thus, we decided to test whether PrP^C^ influences the neuroinvasion of α-syn strains following intraperitoneal (IP) inoculation. M83-*Prnp*^+/+^ and M83-*Prnp*^0/0^ mice were IP-inoculated with M83^+/-^ brain extract containing either the S or NS strain and then monitored longitudinally for the development of motor dysfunction (**Fig. 6a**). As with IC inoculation, there was no significant difference in the kinetics of disease progression for M83-*Prnp*^+/+^ and M83-*Prnp*^0/0^ mice inoculated IP with the S strain (**Fig. 6b**). Opposite to what occurred with IC inoculation, M83-*Prnp*^0/0^ mice developed motor impairment slightly but significantly earlier than M83-*Prnp*^+/+^ mice following IP inoculation. Detergent-insoluble PSyn was present in the brains of both M83-*Prnp*^+/+^ and M83-*Prnp*^0/0^ mice following IP inoculation with either the S or NS strain of α-synuclein aggregates (**Fig. 6c**).

**Figure 6.**
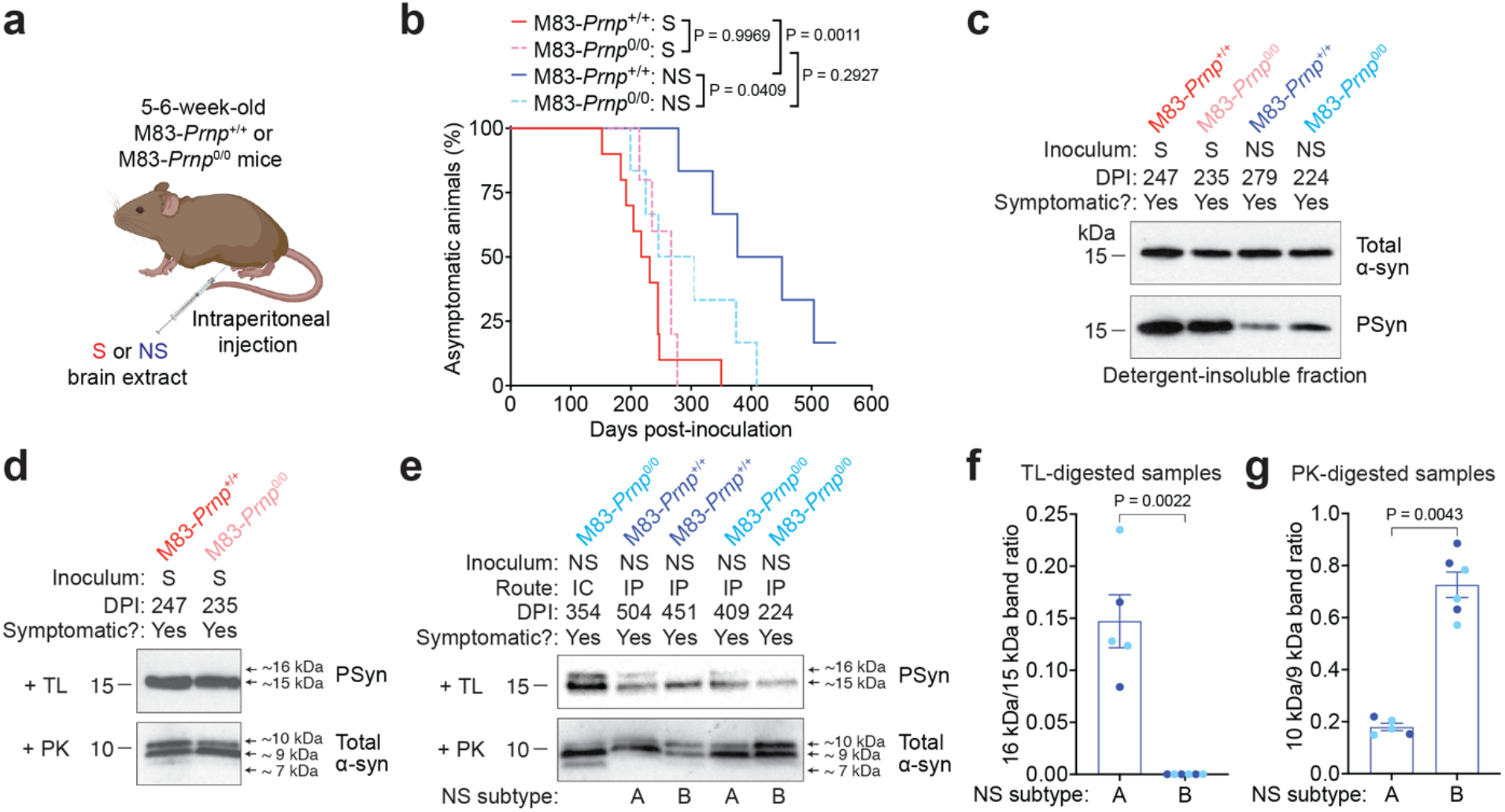
Intraperitoneal inoculation of M83-*Prnp*^+/+^ and M83-*Prnp*^0/0^ mice with the S and NS strains. a) Schematic of intraperitoneal inoculation experiments in M83-*Prnp*^+/+^ and M83-*Prnp*^0/0^ mice. **b)** Kaplan-Meier survival curves for M83-*Prnp*^+/+^ and M83-*Prnp*^0/0^ mice inoculated intraperitoneally with S or NS strain (n = 5-10 mice per experimental condition). Statistical significance was assessed using the Log-rank test. **c)** Immunoblots of detergent-insoluble α-syn in brain extracts from symptomatic M83-*Prnp*^+/+^ and M83-*Prnp*^0/0^ mice at the indicated DPI with S or NS strain, probed for total α-syn and PSyn. **d)** Immunoblots of detergent-insoluble TL-resistant PSyn in brain extracts from symptomatic M83-*Prnp*^+/+^ and M83-*Prnp*^0/0^ mice inoculated intraperitoneally with S strain, as well as detergent-insoluble PK-resistant total α-syn. **e)** Immunoblots for detergent-insoluble TL-resistant PSyn and PK-resistant total α-syn in brain extracts from symptomatic M83-*Prnp*^+/+^ and M83-*Prnp*^0/0^ mice inoculated intraperitoneally with NS strain. Brain extract from an M83-*Prnp*^0/0^ mouse inoculated intracerebrally with NS strain is shown for comparison. Based on banding patterns of protease-resistant α-syn, two different NS subtypes were identified following intraperitoneal inoculation. **f)** Quantification of band intensity ratios in PSyn immunoblots following TL digestion for mice exhibiting NS subtype A (n = 5) or NS subtype B (n = 6). Data for M83-*Prnp*^+/+^ (dark blue) and M83-*Prnp*^0/0^ (light blue) mice is pooled. **g)** Quantification of band intensity ratios in immunoblots for total α-syn following PK digestion for mice exhibiting the two NS subtypes. Data for M83-*Prnp*^+/+^ (dark blue) and M83-*Prnp*^0/0^ (light blue) mice is pooled. The graphs in panels f and g display mean ± s.e.m., and statistical significance was assessed using a Mann-Whitney test.

Like the IC-inoculated mice, the brains of IP-inoculated mice contained α-syn aggregates that were resistant to TL and PK digestion (**Fig. 6d, e**). For both M83-*Prnp*^+/+^ and M83-*Prnp*^0/0^ mice, the pattern of protease-resistant α-syn fragments following IP inoculation with the S strain was very similar to that of mice inoculated IC with the S strain, with a single TL-resistant band at ∼15 kDa and the absence of an ∼7 kDa band following digestion with PK (**Fig. 6d**). Unexpectedly, mice inoculated IP with the NS strain exhibited two distinct patterns of protease-resistant α-syn following digestion of brain extracts with TL or PK. These were termed NS subtype A and subtype B, or NS(A) and NS(B) for short. Following TL digestion, the NS(A) pattern resembled that of mice inoculated IC with the NS strain, with protease-resistant α-syn fragments with molecular weights of ∼15 and ∼16 kDa (**Fig. 6e, f**). In contrast, the brains of mice exhibiting the NS(B) pattern lacked the ∼16 kDa band. Following PK digestion, both NS subtypes exhibited protease-resistant α-syn fragments with molecular weights of ∼10 and ∼9 kDa. However, the ratio of the two bands differed significantly for the two subtypes (**Fig. 6e, g**). It is worth noting that neither NS(A) nor NS(B) completely resembled the pattern present in mice inoculated IC with the NS strain, which is characterized by an additional ∼7 kDa PK-resistant α-syn fragment (**Fig. 6e**). More importantly, the two NS subtypes arose in both M83-*Prnp*^+/+^ and M83-*Prnp*^0/0^ mice that had been inoculated IP with the NS strain. There was no difference in the disease kinetics for mice exhibiting the NS(A) or NS(B) subtypes following IP inoculation (**Fig. S2**).

Both M83-*Prnp*^+/+^ and M83-*Prnp*^0/0^ mice exhibited PSyn deposition in the pons, midbrain, and hypothalamus following IP inoculation with the S or NS strain (**Fig. 7a**). In mice inoculated with the S strain, significantly higher amounts of PSyn pathology were observed in the brains of M83-*Prnp*^+/+^ mice (**Fig. 7b**). However, for the NS strain, no difference in the extent of PSyn deposition was found between M83-*Prnp*^+/+^ and M83-*Prnp*^0/0^ mice following IP inoculation. Interestingly, when the data for NS-inoculated mice was separated into the NS(A) and NS(B) subtypes, as defined by their patterns of protease-resistant α-syn, NS(B) mice exhibited a similar extent of PSyn deposition to S-inoculated mice, whereas NS(A) mice were significantly different than both NS(B) mice and S-inoculated mice (**Fig. 7c**).

**Figure 7.**
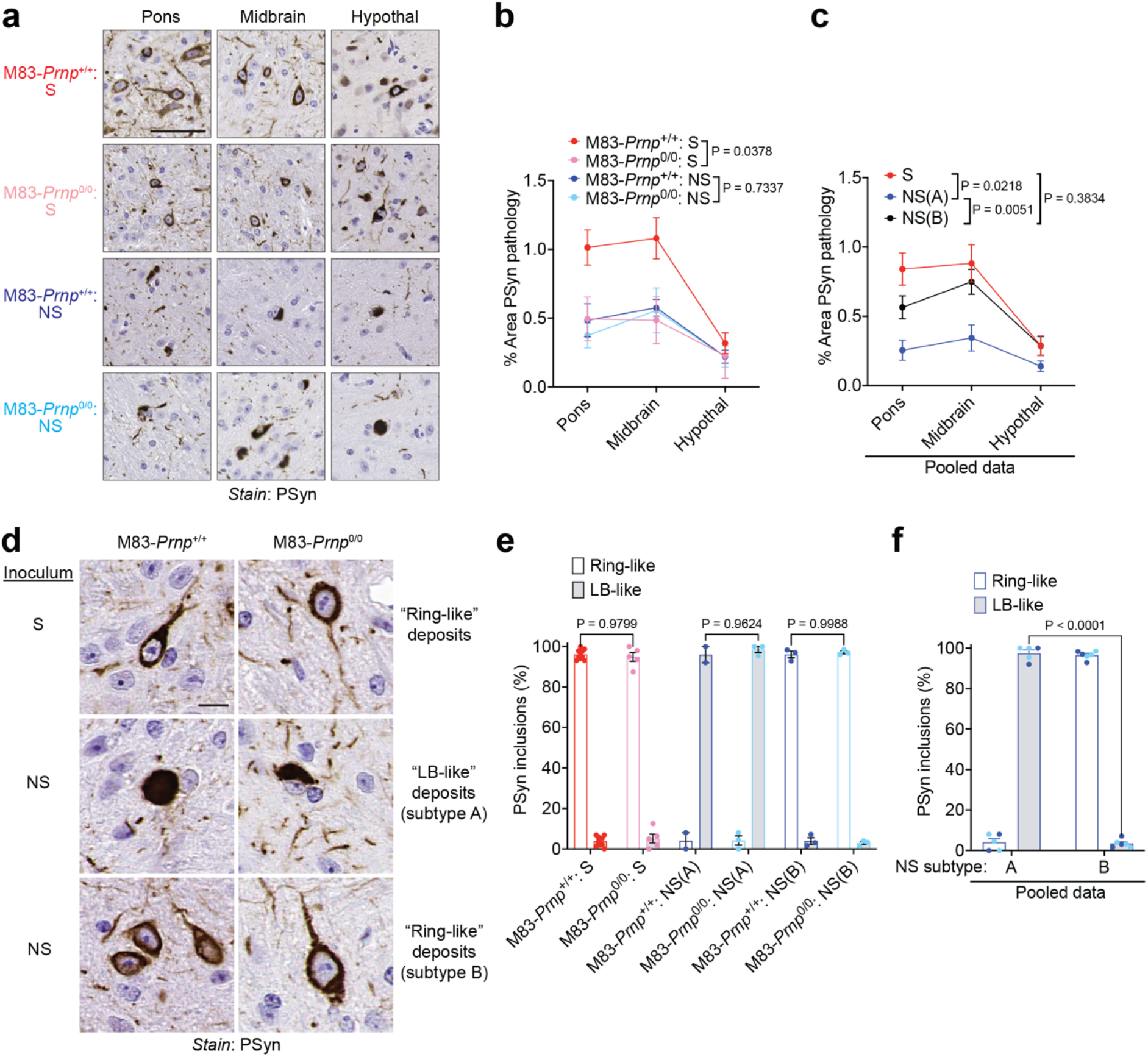
Neuropathology of M83-*Prnp*^+/+^ and M83-*Prnp*^0/0^ mice inoculated intraperitoneally with the S and NS strains. a) Representative images of PSyn-stained sections from the pons, midbrain, and hypothalamus of symptomatic M83-*Prnp*^+/+^ and M83-*Prnp*^0/0^ mice inoculated intraperitoneally with the S or NS strains. Scale bar = 50 µm. **b)** Quantification of the area covered by PSyn staining in sections from the indicated brain regions of symptomatic M83-*Prnp*^+/+^ and M83-*Prnp*^0/0^ mice inoculated intraperitoneally with either S or NS (n = 5-10 mice per experimental condition). **c)** Pooled quantitative PSyn staining data for M83-*Prnp*^+/+^ and M83-*Prnp*^0/0^ mice inoculated intraperitoneally with S strain (n = 15) or exhibiting the NS subtype A (n = 5) or NS subtype B (n = 6) phenotype following intraperitoneal inoculation. **d)** Representative images of PSyn-stained sections from the midbrain of symptomatic M83-*Prnp*^+/+^ and M83-*Prnp*^0/0^ mice inoculated intraperitoneally with the S or NS strains. Scale bar = 10 µm. **e**) Quantification of the relative percentages of ring-like and LB-like PSyn inclusions in the midbrain of S-or NS-inoculated mice (n = 5-10 mice per experimental condition). **f)** Relative percentages of ring-like and LB-like PSyn inclusions in the midbrain for M83-*Prnp*^+/+^ (dark blue) and M83-*Prnp*^0/0^ mice (light blue) exhibiting the NS subtype A (n = 5) or NS subtype B (n = 6) phenotype following intraperitoneal inoculation. The graphs in panels b, c, e, and f display mean ± s.e.m., and statistical significance was assessed using two-way ANOVA.

While the midbrain of both M83-*Prnp*^+/+^ and M83-*Prnp*^0/0^ mice inoculated IP with the S strain contained ring-like PSyn deposits, the morphology of the PSyn aggregates induced by IP inoculation with the NS strain differed by NS subtype (**Fig. 7d-f**). The brains of mice exhibiting the NS(A) phenotype displayed predominantly LB-like PSyn deposits whereas the brains of mice with the NS(B) phenotype contained ring-like deposits. For the S strain as well as the NS(A) and NS(B) subtypes, no significant differences in the morphology of PSyn deposits were observed between M83-*Prnp*^+/+^ and M83-*Prnp*^0/0^ mice following IP inoculation (**Fig. 7e**). However, when data for M83-*Prnp*^+/+^ and M83-*Prnp*^0/0^ mice were pooled, a significant difference was observed between mice exhibiting the NS(A) and NS(B) phenotypes (**Fig. 7f**). Unlike for IC-inoculated mice, the number of cortical PSyn deposits did not differ significantly between mice inoculated IP with the S or NS strains, or between mice exhibiting the NS(A) and NS(B) phenotypes, regardless of *Prnp* genotype (**Fig. S3a, b**). Moreover, hippocampal PSyn deposits were not observed in any of the experimental groups following IP inoculation (**Fig. S3c**), although occasional PSyn-positive astrocytes were selectively present in the thalamus of mice exhibiting the NS(A) phenotype following IP inoculation with the NS strain (**Fig. S3d**). Collectively, these results demonstrate that the NS strain can evolve following IP inoculation into two subtypes: NS(A), which more closely resembles the original NS strain, and NS(B), which is more reminiscent of the S strain. However, both the neuroinvasion and evolution of α-syn strains were largely independent of PrP^C^ expression.

## Discussion

In this study, we examined the propagation and neuroinvasion of two α-syn strains, S and NS, which carry the PD-associated A53T mutation, in M83 transgenic mice that express sequence-matched α-syn. It is now believed that PD can be broken down into two subtypes: “brain-first” PD and “body-first” PD, which differ by the anatomical location in which α-syn aggregates initially arise [71]. The IC injection experiments can therefore be considered as a model of brain-first synucleinopathy, whereas the IP injection experiments constitute a model of body-first synucleinopathy. Given that PrP^C^ has been identified as a putative receptor for α-syn, potentially facilitating the cell-to-cell spread or toxicity of α-syn aggregates [48-50,62], we investigated whether PrP^C^ had any influence on the propagation of α-syn strains in M83 transgenic mice. While PrP^C^ did not modulate the propagation of the S strain following either IC or IP inoculation, the absence of PrP^C^ slightly delayed disease kinetics following IC inoculation with the NS strain. However, the opposite was true following IP inoculation, with M83-*Prnp*^0/0^ mice developing motor impairment faster than M83-*Prnp*^+/+^ mice. We speculate that these conflicting results are more likely to have arisen due to experimental variability than a biologically meaningful effect of *Prnp* genotype. Indeed, the mean incubation period for M83-*Prnp*^0/0^ mice inoculated IC with the NS strain (282 ± 17 days) was comparable to what we previously reported for the NS strain in M83^+/-^ mice (328 ± 21 days), which express PrP^C^ [31]. Given that PrP^C^ was not required for the development of synucleinopathy in any of our experiments and that all other α-syn strain-specific attributes were not substantially modified in the absence of PrP^C^ expression, we conclude that PrP^C^ play a very minor role, if any, in the propagation and neuroinvasion of α-syn strains in mice.

The notion that PrP^C^ can function as a receptor for protein aggregates relevant to human neurodegenerative diseases has been hotly debated. In prion disease, PrP^C^ expression is required for prion neurotoxicity [72,73], and several studies have shown that PrP^C^ facilitates the uptake, spread, and toxicity of α-syn aggregates in cultured cells and mice [48-50,55,62]. However, in line with our findings, another study reported that PrP^C^ is not required for α-syn aggregate-mediated toxicity in neurons or mice [63]. This is reminiscent of the controversy surrounding the role of PrP^C^ in the transduction of neurotoxic signals stemming from Aβ aggregates and its relevance to the pathobiology of Alzheimer’s disease. Many studies have shown that PrP^C^ can bind to oligomeric Aβ aggregates and that Aβ-mediated synaptotoxicity in neurons and mice requires PrP^C^ expression [56,74-77]. In contrast, others have found that Aβ toxicity is independent of PrP^C^ [78-81]. Because there is ample evidence on both sides of the debate and it is well established that PrP^C^ can bind to many different types of protein assemblies [55,57,61,82], it seems likely that PrP^C^’s role as a receptor for protein aggregates may be highly context specific. Thus, while PrP^C^ may be capable of binding α-syn aggregates, this event may not have physiological relevance in all situations. Another possibility is that the spreading and toxicity of α-syn aggregates in M83 mice occurs via a non-receptor-mediated mechanism or involves binding of α-syn assemblies to a different cellular receptor. Extracellular vesicles such as exosomes as well as tunneling nanotubes have been implicated in the cell-to-cell spread of α-syn aggregates [39,41,83], and it is possible that these mechanisms play a larger role in M83 mice or can compensate for the loss of PrP^C^ expression. Alternatively, non-PrP^C^ cellular receptors such as LAG3 could mediate α-syn spread in M83 mice, although it should be noted that, similar to PrP^C^, the involvement of LAG3 in the transmission of α-syn aggregates has been questioned [42,84].

While a definitive explanation for the discrepant results between studies examining the role of PrP^C^ in α-syn aggregate propagation will require further investigation, the specific paradigms used could have a significant impact. We chose to use M83 transgenic mice inoculated with pre-existing α-synuclein aggregates because this model permits the assessment of overt motor dysfunction as well as the effects of different α-syn strains [31,67]. Nevertheless, as with all experimental paradigms, there are caveats and limitations associated with the use of M83 mice. First, as both the S and NS strains as well as M83 mice contain A53T-mutant α-syn, it is plausible that the A53T mutation ablates or diminishes the interaction of α-syn aggregates with PrP^C^. Similarly, mouse PrP^C^ is expressed in the M83-*Prnp*^+/+^ line, and it is possible that the human α-syn aggregates induced by the S and NS strains do not interact as efficiently with mouse PrP^C^ as human PrP^C^. Second, given that small soluble α-syn aggregates such as oligomers have a higher affinity for PrP^C^ than larger fibrils [48,55], α-syn may induce its downstream effects through both PrP^C^-dependent and PrP^C^-independent means, depending on its assembly state. The S and NS strains we used are resistant to treatment with proteases like TL and PK [31], suggesting that they may be composed of larger α-syn aggregates that may not bind as efficiently to PrP^C^ when propagated in M83 mice. Third, the binding between PrP^C^ and α-syn aggregates could be strain-specific. The S and NS strains we utilized exhibit distinct propagation properties in M83 mice than α-syn aggregates from PD patients, implying that structural differences likely exist [67,85]. Finally, it is possible that subtle differences in behavior and/or neuronal physiology existed between the various groups of M83-*Prnp*^+/+^ and M83-*Prnp*^0/0^ mice, which did not exert a major influence on disease kinetics or end-stage neuropathology. However, in agreement with our results, a recent study found no difference in anxiety behavior, nest building, or motor skills between M83^+/-^ mice with and without PrP^C^ expression [62].

Perhaps the most surprising finding of this study was that two disease subtypes emerged in mice that were IP-inoculated with the NS strain, neither of which was an exact phenocopy of the original strain. Approximately half of the mice exhibited the NS(B) phenotype, which was reminiscent of the more rapidly progressive S strain, as revealed by the pattern of protease-resistant α-syn aggregates and the presence of ring-like α-syn deposits in the brain. Interestingly, slowly progressive prion strains can occasionally evolve into a more rapidly progressive strain [86,87]. A similar phenomenon may be occurring in the M83 mice, but the underlying mechanism and why it only occurs following IP inoculation remains to be determined. Two possible explanations seem plausible: 1) during periphery-to-brain propagation of the NS strain, conformational templating may not be completely faithful to the strain’s original structure, leading to strain mutation and the emergence of a new variant that propagates faster and thus has a selective advantage; and 2) the NS strain may exist as a “cloud” of aggregate conformations, and IP inoculation results in the selection of an S-like minor species that can more readily propagate to the brain [88]. The generalizability of this finding to other α-syn strains, including those derived from human brains, warrants further investigation. Given that the NS(A) and NS(B) subtypes were observed equally frequently in both M83-*Prnp*^+/+^ and M83-*Prnp*^0/0^ mice, the presence or absence of PrP^C^ had no influence on α-syn strain evolution.

Our findings in M83 transgenic mice suggest that PrP^C^ may not be an ideal therapeutic target for blocking the cell-to-cell propagation of α-syn aggregates in synucleinopathies such as PD and MSA. Antibodies against PrP^C^ prevent the accumulation of prions in cultured cells and mice and, despite limited testing so far, have shown promise for the treatment of human prion disease [89-93]. Moreover, anti-PrP antibodies that block the interaction between PrP^C^ and Aβ aggregates attenuate toxicity and cognitive deficits in rodents and thus are being considered for use as an Alzheimer’s disease therapeutic [94-97]. On the other hand, certain anti-PrP antibodies can elicit neurotoxicity [98-100]. While it is conceivable that immunotherapy against PrP^C^ may have subtle beneficial effects that could be used in combination with other treatment modalities, anti-PrP antibodies on their own may not significantly interfere with disease progression in the synucleinopathies.

## Acknowledgements

The authors thank Gabor Kovacs and Ali Karakani for assistance with the scanning of pathology slides. This work was supported by grants to JCW from the Scottish Rite Charitable Foundation of Canada (#18119) and the Canadian Institutes of Health Research (PJT-169042). RWLS was supported by fellowships from the Croucher Foundation, Parkinson Canada, the Peterborough K.M. Hunter Charitable Foundation, as well as an Ontario Graduate Scholarship. Experimental schematics were created using BioRender.com.

## Declarations

### Availability of data and material

All data generated or analyzed during this study are included in the manuscript.

### Author contributions

RWLS and JCW contributed to the study conception and design. Material preparation, data collection and analysis were performed by RWLS, ES, and AEA. AA and GLC contributed novel experimental tools. The first draft of the manuscript was written by RWLS and JCW. All authors read and approved the final manuscript.

