## Supplemental information for "α-Synuclein Strain Propagation is Independent of Cellular Prion Protein Expression in Transgenic Mice"

### Supplemental Table 1. List of mice with intercurrent illness removed from the study

| Inoculum, route | Mouse genotype | Animal ID | Sex | Days post inoculation | Notes | Protease digestion pattern |
|---|---|---|---|---|---|---|
| S, IC | M83-*Prnp*$^{+/+}$ | 7790 | F | 93 | Euthanized due to atypical illness | --- |
| | | 7801 | F | 166 | Euthanized due to atypical illness | --- |
| | | 7802 | M | 100 | Euthanized due to fight wounds | --- |
| | | 7803 | M | 152 | Found dead | Brain not collected |
| S, IC | M83-*Prnp*$^{0/0}$ | 7665 | F | 163 | Euthanized due to atypical illness | --- |
| | | 7681 | F | 166 | Euthanized due to atypical illness | --- |
| | | 6676 | M | 195 | Euthanized due to fight wounds and urinary blockage | S |
| | | 6677 | M | 166 | Found dead due to fighting injuries | Brain not collected |
| | | 6673 | M | 166 | Found dead due to fighting injuries | Brain not collected |
| | | 7671 | M | 145 | Euthanized due to atypical illness | --- |
| NS, IC | M83-*Prnp*$^{+/+}$ | 7554 | M | 192 | Euthanized due to fight wounds | NS |
| S, IP | M83-*Prnp*$^{+/+}$ | 9851 | F | 196 | Euthanized due to atypical illness | --- |
| | | 8868 | M | 156 | Euthanized due to fight wounds | --- |
| | | 8878 | M | 152 | Found dead | --- |
| | | 9858 | M | 231 | Euthanized due to atypical illness | --- |
| S, IP | M83-*Prnp*$^{0/0}$ | 9056 | F | 116 | Found dead | Brain not collected |
| | | 8737 | M | 270 | Euthanized due to atypical illness | --- |
| NS, IP | M83-*Prnp*$^{+/+}$ | 8493 | F | 171 | Euthanized due to atypical illness | --- |
| | | 9394 | F | 370 | Euthanized due to atypical illness | --- |
| | | 8500 | M | 157 | Found dead | Brain not collected |
| | | 8502 | M | 168 | Found dead | Brain not collected |
| NS, IP | M83-*Prnp*$^{0/0}$ | 8740 | F | 375 | Euthanized due to atypical illness | --- |
| | | 8482 | M | 192 | Euthanized due to atypical illness | --- |
| | | 8484 | M | 227 | Euthanized due to atypical illness | --- |
| | | 8485 | M | 227 | Euthanized due to atypical illness | --- |
| | | 8731 | M | 109 | Euthanized due to atypical illness | --- |
| | | 8739 | M | 160 | Found dead | Brain not collected |

Three dashes (---): Brains collected and analyzed by protease digestion assay; no protease-resistant α-syn detected by immunoblotting.

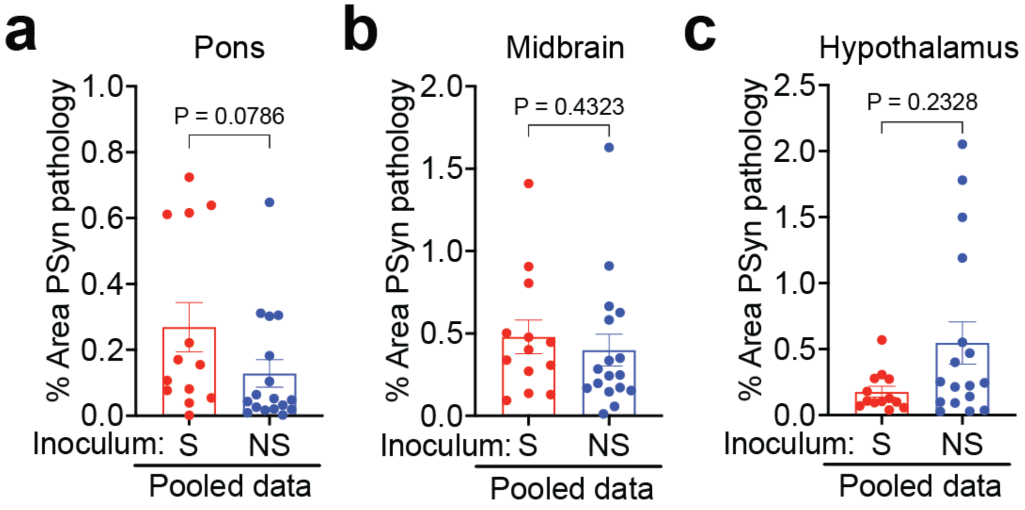


**Supplemental Figure 1. The extent of PSyn deposition in the hindbrain does not differ between M83 mice inoculated intracerebrally with the S or NS strains.** Pooled quantitative data for the area covered by PSyn staining in the pons (**a**), midbrain (**b**), and hypothalamus (**c**) of symptomatic M83-*Prnp*$^{+/+}$ and M83-*Prnp*$^{0/0}$ mice inoculated intracerebrally with either S (n = 13) or NS (n = 17) strain. The graphs display mean ± s.e.m. Statistical significance was assessed using a Mann-Whitney test.

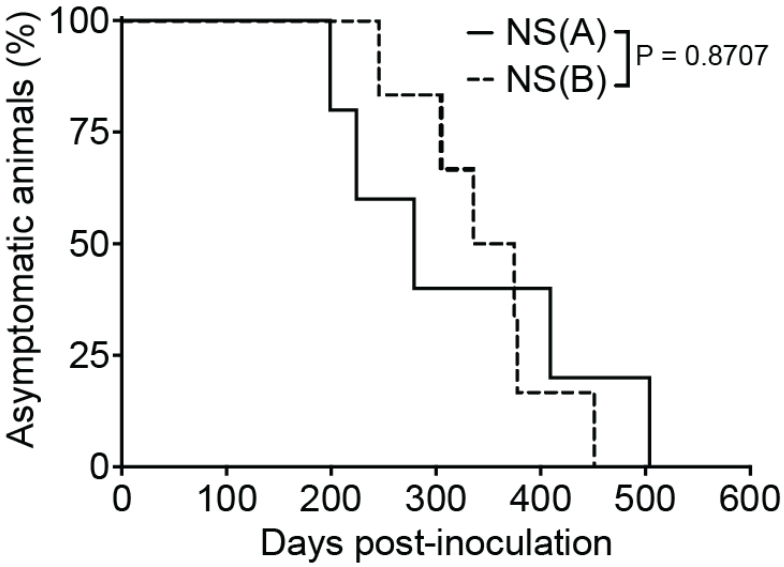


**Supplemental Figure 2. The NS(A) and NS(B) subtypes do not exhibit distinct incubation periods.** Kaplan-Meier survival curves for M83-*Prnp*$^{+/+}$ and M83-*Prnp*$^{0/0}$ mice (pooled data) inoculated intraperitoneally with the NS strain. Based on patterns of protease-resistant α-syn, mice were classified as either subtype A ($n = 5$) or B ($n = 6$). Statistical significance was assessed using the Log-rank test.

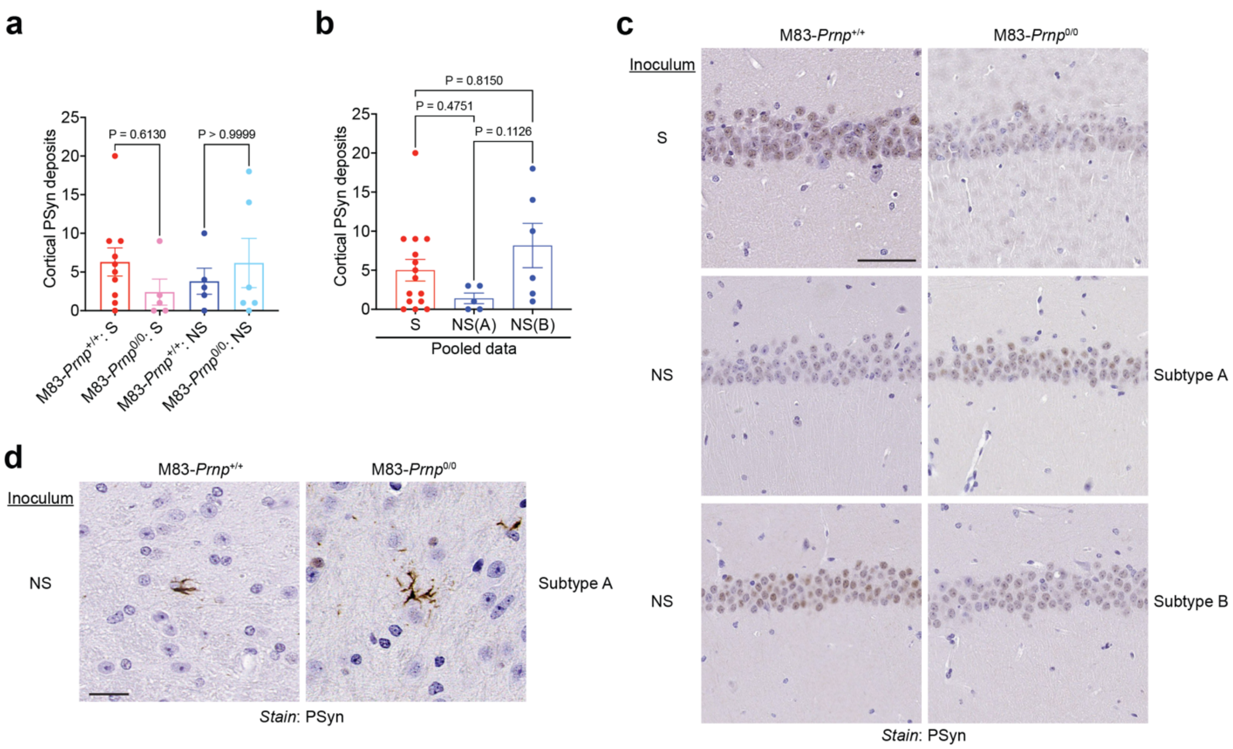


**Supplemental Figure 3. Evaluation of α-syn strain-specific brain region and cell type targeting in intraperitoneally inoculated M83-*Prnp*$^{+/+}$ and M83-*Prnp*$^{0/0}$ mice. a)** Quantification of the number of PSyn-positive neurons in the cortex of symptomatic M83-*Prnp*$^{+/+}$ and M83-*Prnp*$^{0/0}$ mice inoculated intraperitoneally with the S or NS strain (n = 5-10 mice per experimental condition). **b)** Pooled PSyn-positive neuronal counts in the cortex of M83-*Prnp*$^{+/+}$ and M83-*Prnp*$^{0/0}$ mice inoculated intraperitoneally with S strain (n = 15) or exhibiting the subtype A (n = 5) or subtype B (n = 6) phenotype following intraperitoneal inoculation with the NS strain. The graphs in panels a and b display mean ± s.e.m., and statistical significance was assessed using a Kruskal-Wallis test followed by Dunn's multiple comparisons test. **c)** Representative images of PSyn-stained sections from the hippocampal CA1 region of symptomatic M83-*Prnp*$^{+/+}$ and M83-*Prnp*$^{0/0}$ mice inoculated intraperitoneally with the S or NS strains. Scale bar = 50 µm. **d)** Representative images of PSyn-stained sections from the thalamus of symptomatic M83-*Prnp*$^{+/+}$ and M83-*Prnp*$^{0/0}$ mice inoculated intraperitoneally with the NS strain and exhibiting the subtype A phenotype. Scale bar = 20 µm.
